# Rate of *de novo* mutations in the three-spined stickleback

**DOI:** 10.1101/2024.11.13.623523

**Authors:** Chaowei Zhang, Kerry Reid, Mikkel Heide Schierup, Hongbo Wang, Ulrika Candolin, Juha Merilä

## Abstract

Being a fundamentally important genetic parameter and evolutionary force, estimates of germline mutation rates have many uses in evolutionary biology. However, accurate estimates of *de novo* mutation (DNM) rates are still relatively scarce even for extensively studied evolutionary biology models. We estimated DNM rates for the three-spined stickleback (*Gasterosteus aculeatus*), the ‘supermodel’ of ecology and evolutionary biology. Using a large number of family trios sequenced to depth of 45x coverage, we identified 115 unique mutations genome wide and estimated the DNM rate at *µ* = 5.05 × 10^-9^/bp/gen without any detectable sex bias. The localised DNM rate was found to be positively correlated with recombination rate supporting the notion that recombination is a mutagenic process. Comparison of *µ* and genomic characteristics to those of the related nine-spined stickleback (*Pungitius pungitius*) revealed a high degree of similarity suggesting that despite 17.5 million years of independent evolution, the mutational processes in the two species appear to have been conserved.

## Introduction

The *de novo* mutation rate, *µ*, is the ultimate source of new genetic variation and a central parameter for a lot of population and evolutionary genetic inference. However, until recently, direct estimates of *µ* were hard to obtain, but the rapid development and refinement of modern sequencing technologies and computational tools have led to an increase in *µ* estimates from various non-model species (Smed, et al. 2016; Koch et al. 2019; Wang, et al. 2022; Sendell-Price et al. 2023; Wang and Obbard 2023; Zhang et al. 2023). Yet, there are still strong taxonomic biases in the available estimates with mammalian, and those of primates in particular, estimates being overrepresented, whereas those of birds, reptiles, amphibians and fish are still rare (Wang and Obbard 2023; but see: Bergeron et al. 2023). Furthermore, there is a clear bias towards estimates from well-established model species (e.g. *H. sapiens, M. musculus, D. melanogaster, C.elegans, Z. mays, A. thaliana*) being based on a much larger number of mutations than those from non-model species. Consequently, most of the *µ* estimates for the non-model species are based on only a few parent-offspring trios and therefore have broad confidence intervals covering several fold ranges of *µ*-values (e.g. Wang and Obbard 2023 their Figure 2; Bergeron et al. 2023 their Figure 1a). Uncertainty around these estimates is further compounded by difficulties controlling false positive and negative rates when estimating *µ* from parent-offspring data (Yoder and Tiley 2021; Bergeron et al. 2022). Hence, there is a definitive need to obtain more accurate estimates of *µ* from many non-model species based on larger numbers of trios.

**Figure 1.**
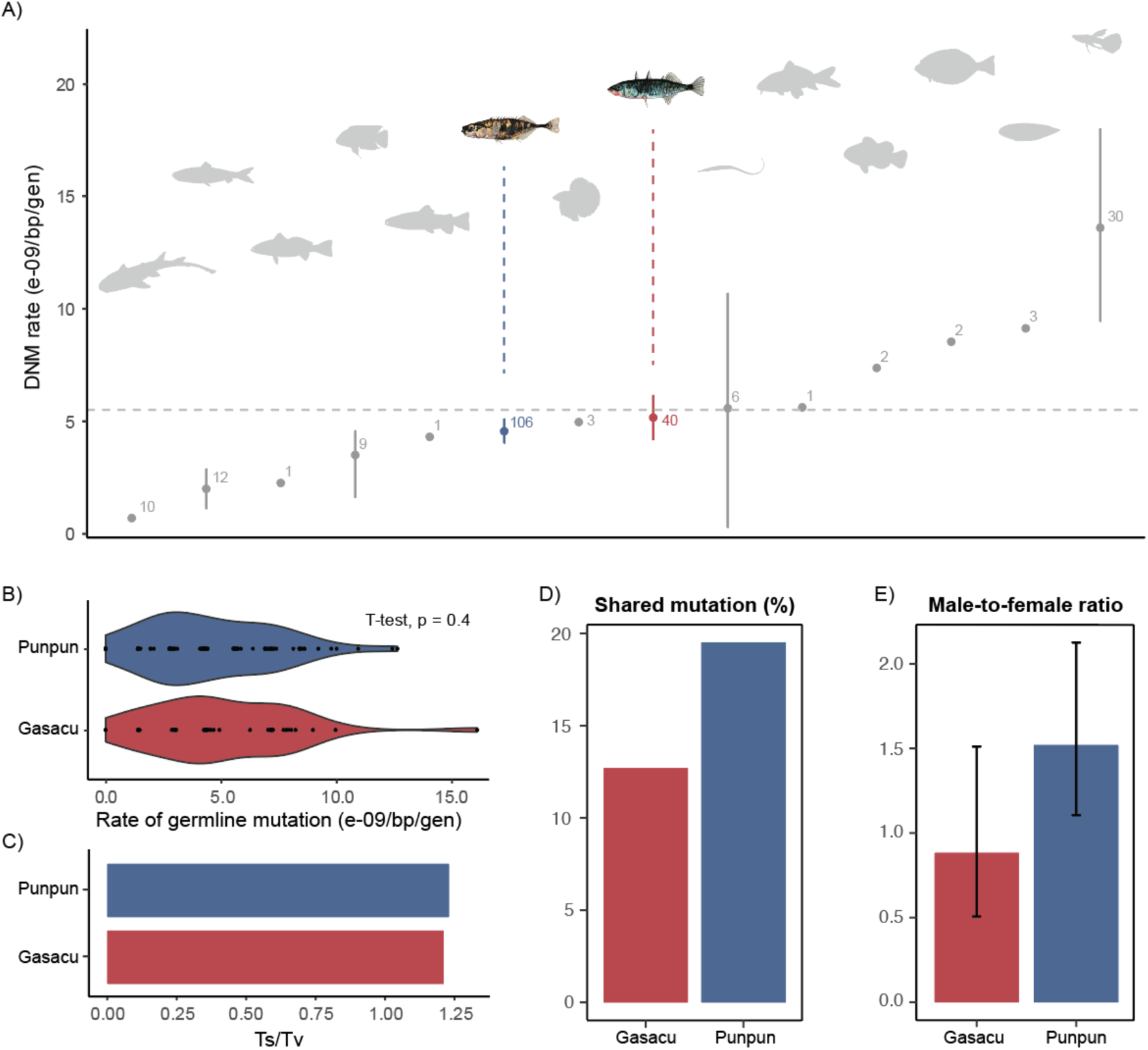
Germline mutation rates (µ) and characteristics in three- (Gasacu) and nine-spined sticklebacks (Punpun). (A) Comparison of *µ in* sticklebacks with those in other fishes, (B) a violin plot of *de novo* mutation (DNM) rates, (C) The transition (Ts) to transversion (Tv) ratio, (D) the percentage of shared mutations among full sibs, and (E) the parental origin (male-to-female ratio) of the DNMs in three- (red) and nine-spined (blue) sticklebacks.

**Figure 2.**
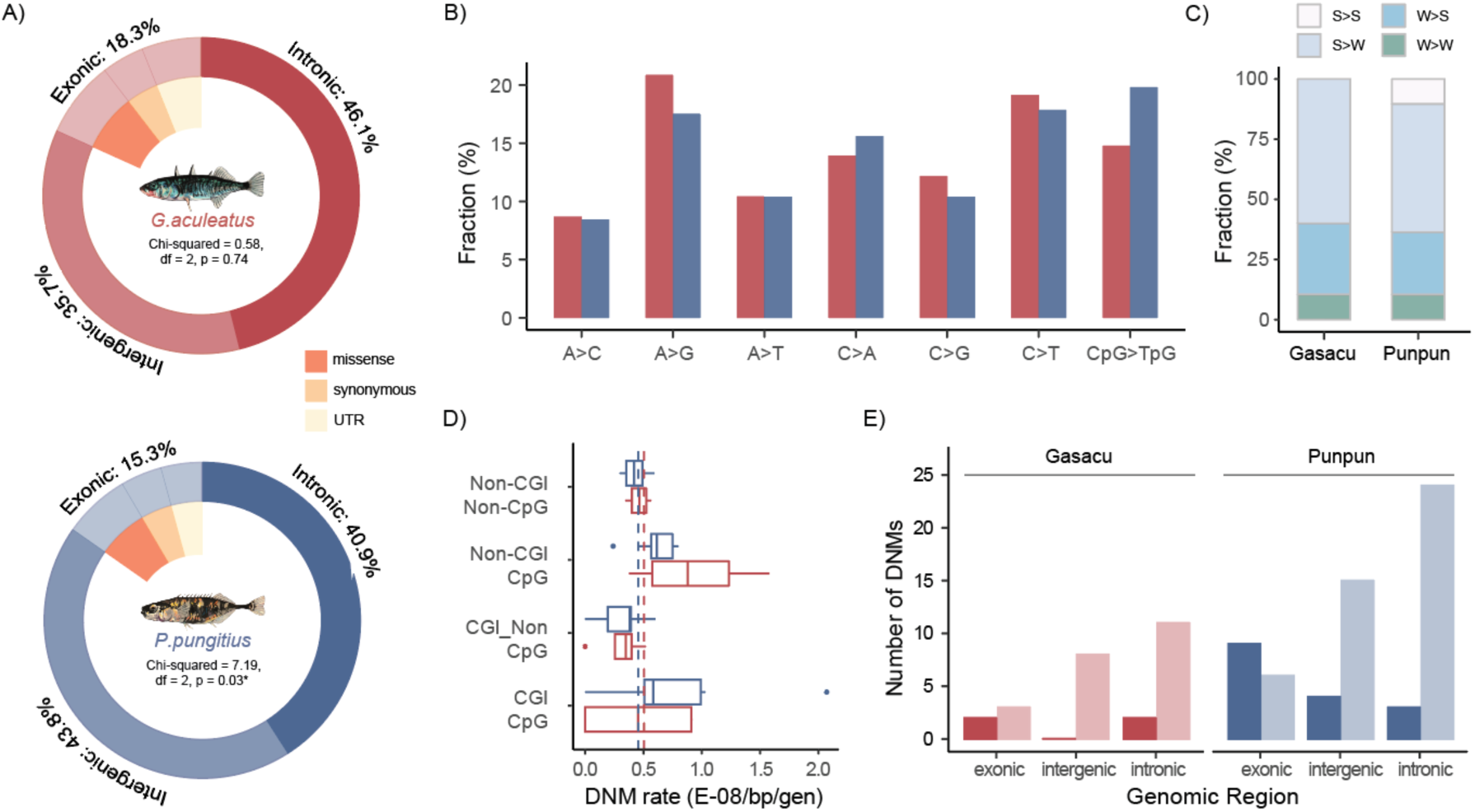
Genomic landscape of de novo mutations (DNMs) in sticklebacks. (A) Distribution of DNMs on different genomic regions. (B) Comparison of mutation spectra between three- (red) and nine- (blue) spined sticklebacks. (C) DNM composition based on changes in pairing types. (D) Pedigree-level DNM rates in CpG or Non-CpG sites within or outside CpG islands (CGI). The two dashed lines depict genomewide DNM rates for the two species. (E) Number of unique CpG DNMs located inside CGI (darker bars) or outside CGI (lighter bars) in the two species.

The three-spined stickleback (*Gasterosteus aculeatus*) is a popular model system for ecological (Bell and Foster 1994), evolutionary (Colosimo et al. 2005; Lescak et al. 2015; Wootton 2009), behavioural (Von Hippel 2010; Bell et al. 2016; Bell et al. 2018; Norton and Gutierrez 2019), physiological (Von Hippel et al. 2018) and genomic (Peichel and Marques 2017; Reid et al. 2021) research and it has been dubbed the “supermodel of ecology and evolutionary biology” (Gibson 2005; Barber 2010). Despite the voluminous body of research tracking its evolutionary history in different parts of this distribution range (Orti et al. 1994; Mäkinen et al 2006; Mäkinen and Merilä 2008; Shikano et al. 2010; Lescak et al. 2015; Fang et al. 2020a,b; Fang et al. 2021; Feng et al. 2022) and research devoted to deciphering genomic underpinnings of its adaptation to different environments (Jones et al. 2012; Fang et al. 2021; Roberts-Kingman et al. 2021), no study has yet estimated its germline *de novo* mutation rates. Studies which have needed to parametrise three-spined stickleback mutation rate have used either substitution rates or generic ball-park estimates of *µ* (Liu et al. 2018; Ravinet et al. 2018; Varadharjan et al. 2019; Yamasaki et al. 2020; Dahms et al. 2022; Feng et al. 2022).

The aim of this study was to estimate the rate of germline *de novo* mutations in the *G. aculeatus* and characterise its mutation spectrum using a large number of family trios sequenced to deep coverage of depth. To gain insights into the evolutionary conservation of mutational processes, we compared the obtained estimates with those of its relative, the nine-spined stickleback (*Pungitius pungitius*), obtained and analysed in a comparable manner.

## Material and Methods

### Sampling and breeding

The fish used in this study originated from the Baltic Sea site Tvärminne, Finland (59°50’20”N; 23°12’15”E). The parental fish were caught with a beach-seine in May 2023 and brought alive to the Tvärminne Zoological Station’s aquaculture facilities. Randomly selected males in breeding condition, as determined by their nuptial coloration, were placed into individual 80L flow-through aquaria with nesting materials to enable nest building (Candolin 1997). When a male had built a nest, a gravid female was placed into the aquarium and allowed to spawn in the nest. If the female did not spawn within 2 hours, she was replaced by another gravid female. After spawning, the female was removed and the male was left to care for the eggs in the nest. When the eggs started to hatch and the first fry appeared outside the nest, the male was removed to prevent him from cannibalising the fry. The fry were fed *Artemia salina* larvae and allowed to grow until 1 week old. All individuals of a family (male, female and offspring) were preserved in 91% ethanol and stored at −20°C before being preserved in 95% alcohol and shipped to Hong Kong for DNA extractions.

### DNA extraction and sequencing

DNA was extracted from the fin clips using the DNEasy Blood and Tissue kit (Qiagen, Germany) with RNAse treatment following the manufacturer’s specifications. After evaluating the DNA purity and the concentration, the DNA samples were sent for PCR-free library preparation and whole genome resequencing to a target depth of 45x coverage (PE150) using the DNBseq platform (Berry Genomics, China).

### Read mapping, variant calling and pedigree examination

The paired-end raw reads were mapped to the reference genome of three-spined stickleback (version 5, https://stickleback.genetics.uga.edu/downloadData/, an updated version to Peichel et al. 2020), using BWA-MEM (v0.7.17, Li and Durbin 2009). The paired reads were sorted, and the mate information was filled in by applying the ‘sort’ and ‘fixmate’ functions in SAMtools. (v1.16.1, Li et al. 2009). The duplicate reads were marked through PicardTools (v2.27.5; http://picard.sourceforge.net). The single nucleotide variants were called for each sample separately from these sorted BAM files using GATK (v4.3.0.0; Van der Auwera and O’Connor 2020) HaplotypeCaller in ERC mode, following the best practices workflow of GATK (Poplin et al. 2017). The resulting gVCF files were then combined and jointly genotyped using GATK CombineGVCFs and GenotypeGVCFs modules for all samples involved. Utilising the hard-filtered output sets of SNPs and indels from this step, Base-quality score recalibration (BQSR) was carried out to the sorted BAM files in GATK, after which the variants were called again from the final BAM files for each individual, and the generated gVCF files were combined in the same manner as mentioned for each parent-progeny trio.

Before the further steps, the parent-offspring relationships within each pedigree were confirmed by estimating the probabilities of identity-by-descent (IBD) with PLINK (v1.90; Chang et al. 2015). The ratio of probabilities of sharing 0, 1 and 2 alleles (Z_0_:Z_1_:Z_2_) between a parent and its offspring was expected to be close to 0:1:0.

### Identifying putative de novo mutations

For each parent-offspring trio, only SNPs violating Mendelian expectations were considered as candidate point mutations. In other words, one of the two alleles of the offspring should be absent from its parents to count as a mutation. Given the rarity of alternative alleles mutating back to reference alleles, we retained only those variants where the parents were homozygous for the reference allele (0/0), and the offspring were heterozygous (0/1). The site filters (also known as the hard filtering suggested by GATK best pipeline, Poplin et al. 2017) and the individual filters were applied following the “Mutationathon” guidelines (Bergeron et al. 2022), as detailed in Supplementary Table 1 and Zhang et al. (2023).

To further control the false positive rates, the putative DNMs were checked by loading the final alignments of each trio in the visualisation tool, IGV (Thorvaldsdóttir et al. 2013), and only sites where the genotypes were consistently supported by the aligned reads were retained. Additionally, the number of reads at the DNM candidates was double-checked in ‘bam-readcount’ (Khanna et al. 2022) to exclude mismatches between the final BAM files and the realigned VCF files. In the end, we retained only those sites where the parents carry pure reference alleles, and the offspring were confirmed to be real heterozygotes after excluding the incorrectly aligned reads.

### De novo mutation rate estimation

The per-generation point mutation rates was estimated for each sample by:

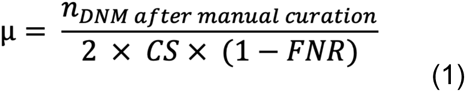

And for all samples by a zero-inflated method:

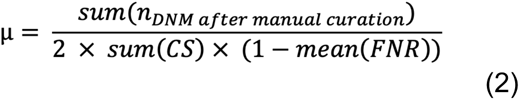

where the callable genome size (CS) was the number of sites along the whole genome for each offspring that passed the depth filtering within its trio family (0.5DP_trio_ < DP_child_ < 2DP_trio_). The false discovery rate (FDR) was the proportion of DNM candidates removed by the IGV and the bamcount inspections mentioned in the above section:

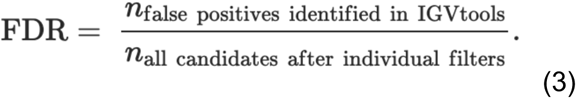

The false negative rate (FNR) was inferred from the percentage of true offspring heterozygotes, when the genotypes of the parental sites were one 0/0 and one 1/1, that had been deleted by the allelic balance (AB) filtering (AB < 0.3 and AB > 0.7).

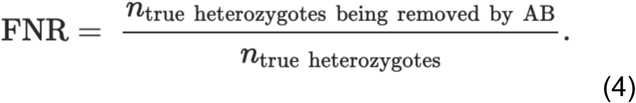

### Genome-wide mutation rate and its comparison with recombination rate

We calculated the genome-wide mutation and recombination rate for the entire dataset using a 5Mb sliding window and a 1Mb step length. Applying the linkage map available for three-spined stickleback (Kivikoski et al. 2022), the recombination fractions were calculated with the following equation:

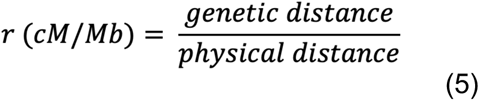

Since recombination is known to be mutagenic, we tested for correlation between mutation and recombination rates. To align with the linkage map, the detected DNMs were lifted over to the corresponding version of the reference genome (Peichel et al. 2017) by utilising the provided chain file (https://stickleback.genetics.uga.edu/downloadData/). We then estimated the per-5Mb DNM count for the same sliding windows. Pearson product moment correlation tests were conducted to see if the per-window size mutation rate correlates with recombination rate.

### Mutation spectrum and genomic context

The CpG islands (Henceforth: CGIs), which are the regions of significantly higher levels of CpG dinucleotides compared to the genome as a whole, were predicted with the default settings using the “twoBitToFa” module and the cpg_lh script in the UCSC tools (http://genome.ucsc.edu/cgi-bin/hgTrackUi?g=cpgIslandExt, Miklem and Hillier, unpublished). The detected CGIs were further classified into intergenic, intronic, exonic, transcription start site (TSS), and transcription termination site (TTS) CGIs, based on their genomic location. With the above statistics, the per-5Mb CpG densities and gene densities were quantified similarly to what was done for the genome-wide mutation/recombination frequencies.

The detected DNMs were classified using both snpEff (v5.1f) and a manual verification according to the annotation file. To generate the mutation spectrum, these mutations were also stratified into six groups according to their mutation forms (A > C, A > T, A > G, C > A, C > G, C > T), with a special inspection on the C > T mutations on the CpG sites (the 7^th^ group). Among these mutation types, A > G and C > T are the transitions (Ts), and the rest are transversions (Tv). A Chi-squared test was used to test whether there was any bias in strong base pairs (C:G) mutating to weak ones (A:T), or vice versa.

To examine if *de novo* mutations were biased to originate from males or females, all the identified DNMs were phased back to their parent of origins with a read-backed phasing method, POOHA (https://github.com/besenbacher/POOHA). The per-sample percentage of the paternal sites was then calculated to infer the paternal-to-maternal ratio (alpha) of the DNMs. Additionally, mutations occurring at the early stages of meiosis and shared among the full-sibs were also identified and counted.

### Comparison to nine-spined stickleback (Pungitius pungitius)

To compare the three-spined stickleback mutation rates and patterns with those of the nine-spined stickleback, we estimated the per-5Mb genome-wide mutation frequencies, recombination frequencies, CpG densities, and the gene densities for *P. pungitius* using exactly the same methods described in the above sections using data from Zhang et al. (2023). The pairwise synteny between the two reference genomes (*G. aculeatus* V4 vs. *P. pungitius* V7) was visualised using MCscan (Tang et al. 2024). The recombination fraction was obtained from a previously available linkage map of the eastern European lineage of *P. pungitius* (Kivikoski et al. 2022). To make the annotations of *P. pungitius* comparable to the three-spined stickleback, we again lifted over the NCBI annotations for the version 6 reference genome (GCF_902500615.1) to the version of the published mutation data and the linkage map (version 7, GCA_902500615.3, Kivikoski et al. 2021) with removing the non-protein-coding genes by the pygtftk tool (Lopez et al. 2019). In total, more than 20 thousand genes were annotated across the autosomal genomes for both species, and the genomes were subsequently divided into exonic (including protein-coding regions and untranslated regions, UTRs), intronic, and intergenic regions.

*t*-tests were conducted to assess the significance of the differences between the two species. To investigate the relationship between various factors and mutation rates, generalised mixed linear models (using a poisson distribution) were constructed in lme4 (Bates et al. 2015) using combined data from the two species. In these models, species nested with the pedigree was treated as a random effect, while other factors were treated as fixed effects:

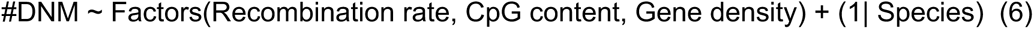

## Results

### De novo mutation rate in three-spined sticklebacks

48 samples belonging to four pedigrees of unrelated parents, each with 10 full-sib offspring, were whole-genome resequenced. The final bam files from the raw reads mapped to the reference genome had a mean coverage of depth of 45.4× before variant calling. Over 526,000 sites were retained across the 40 trios after filtering for Mendelian violations in all autosomal variants (excluding those on the sex chromosomes Y and XIX), averaging 13,152 sites per progeny. After application of a series of filters, a total of 115 unique DNMs were detected (see Supplementary Table 1 for more details). The average callable genome size was 87.40% of the entire autosomal genome (363.45Mb), and the mean false negative rate (FNR) was 7.44%. The overall average per-generation point DNM rate was estimated to be *µ* = 5.05 × 10^-9^/bp/gen (95% confidence interval: 4.04 – 6.07 × 10^-9^/bp/gen, with zero-inflated method being 5.02 × 10^-9^/bp/gen), and it resided in the middle of all current available DNM rates in fish, with the second narrowest 95% CIs of all fish species (Figure 1A). Assuming a generation time of three years for the marine three-spined sticklebacks, the yearly mutation rate would be on average 1.68 × 10^-9^/bp/year. The per-generation DNM rates did not differ between marine three- and nine-spined sticklebacks (Figure 1B, t_64.44_ = 0.85; p = 0.40).

### Mutation characteristics and spectrum

Out of all distinct DNMs identified, 54.8% were transitions (Ts) and 45.2% were transversions (Tv), resulting in a Ts:Tv ratio of 1.21, which was similar to that in the nine-spined stickleback (Figure 1C). In both species, we observed about twice as many DNMs mutating from the strong (C:G) to weak (A:T) base pair (S>W) compared to the opposite direction (W>S), and very few DNMs mutating within its own pairing type (S>S or W>W, Figure 2C). This finding deviates significantly from the null expectation of equal mutation rates, given the GC proportion (*G. aculeatus*: χ^2^ = 18.30, df = 1, p = 1.88e-05; *P. pungitius*: χ^2^ = 50.57, df = 1, p = 1.15e-12). This GC mutation bias was also evident in the mutation spectrum (Figure 2B). In addition to the typically high proportion of C>T mutations, whether at CpG sites or not, an elevated number of C>A mutations was observed in both species as well (Figure 2B).

In total, 21, 53 and 41 DNMs were found in the exonic, intronic, and intergenic regions in *G. aculeatus*, while these numbers were 47, 126 and 135 in *P. pungitius*, respectively. The DNM locations did not deviate significantly from a random distribution across genomic features in *G. aculeatus* (χ^2^ = 0.58, df = 2, p = 0.74), but slightly so in *P. pungitius* with fewer DNMs detected within the intronic regions than expected (χ^2^ = 7.19, df = 2, p = 0.03, Figure 2A). Among the exonic mutations for the two species, 45.6% were missense, 23.5% were synonymous DNMs and 30.9% occurred on UTRs (Figure 2A). These exonic mutations were all categorised as being either moderate or low impact by snpEff, and no loss-of-function mutations were observed in either dataset.

The total length of the autosomal CGI regions in three-spined sticklebacks was estimated to be 11.8% of the entire genome. 9.6% of the unique DNMs were located within these areas, with five DNMs being C>T and three situated on CpG sites. There was a clear inflation in DNM rates in the non-CGI CpG sites in three-spined sticklebacks (Figure 2D), although this was not significantly different from a null expectation given the GC contents of the CGI or non-CGI regions (χ^2^ = 0.15, df = 1, p = 0.70). These CGI CpG DNMs were located only on exons and the introns (χ^2^ = 24.33, df = 2, p = 5.2e-06; the pink bars in Figure 2E), while the non-CGI ones were located randomly in respect to genomic locations (χ^2^ = 2.35, df = 2, p = 0.31; the red bars in Figure 2E). The pattern was similar in the nine-spined sticklebacks as demonstrated by the light and dark blue bars in Figure 2E.

### Weak sex bias of DNM rates in sticklebacks

In total, 64.3% unique DNMs detected in the offspring were traced back to their parents-of-origin, among which on average 46.9% (95% CI: 33.6% – 60.2%) DNMs originated from the fathers. This translated to an α value (male-to-female ratio) at 0.88 (95% CI: 0.51 – 1.51), which was not significantly different from that of the nine-spined stickleback (Figure 1E, t_61.43_ = −1.75, p = 0.085). Unlike in *P. pungitius* in which a significantly higher proportion of CpG sites were inherited from the paternal than the maternal side (Supplementary Figure 1), the proportions were equal in the three-spined stickleback (χ^2^ = 0.22, df = 1, p = 0.64).

Additionally, less DNMs were found to be shared among full-sibs in the three-spined than in the nine-spined sticklebacks (12.7% *vs.* 19.5%, Figure 1D). Except for DNM shared between two offspring, sites shared between 3 and 4 siblings were observed on rare occasions. Due to the limited number of shared mutation being phased back to their parental origins (7 and 48 in three- and nine-spined sticklebacks, respectively), a combined dataset of the two species were analysed, and no sex bias was detected neither for the shared (Maternal:Paternal = 26:29, χ^2^ = 0.16, df = 1, p = 0.27) nor for the non-shared DNMs (Maternal:Paternal = 108:125, χ^2^ = 1.24, df = 1, p = 0.69).

### Mutation distributions and their correlation with the other genomic features

The synteny analyses demonstrated similar trends between the two species in their genome-wide features, including the mutation frequencies, recombination rates, the CpG dinucleotide contents and the gene densities (Figure 3A). Notably, for the translocation identified between chr7 (1-13.2Mb) in *G. aculeatus* and the pseudoautosomal region on LG12 (from 16.9Mb) in *P. pungitius* (Figure 3A), the *de novo* mutation rates were similarly high exceeding the genome-wide level, at respectively 1.10 × 10^-8^/bp/gen and 9.94 × 10^-9^/bp/gen (Supplementary Figure 3B).

**Figure 3.**
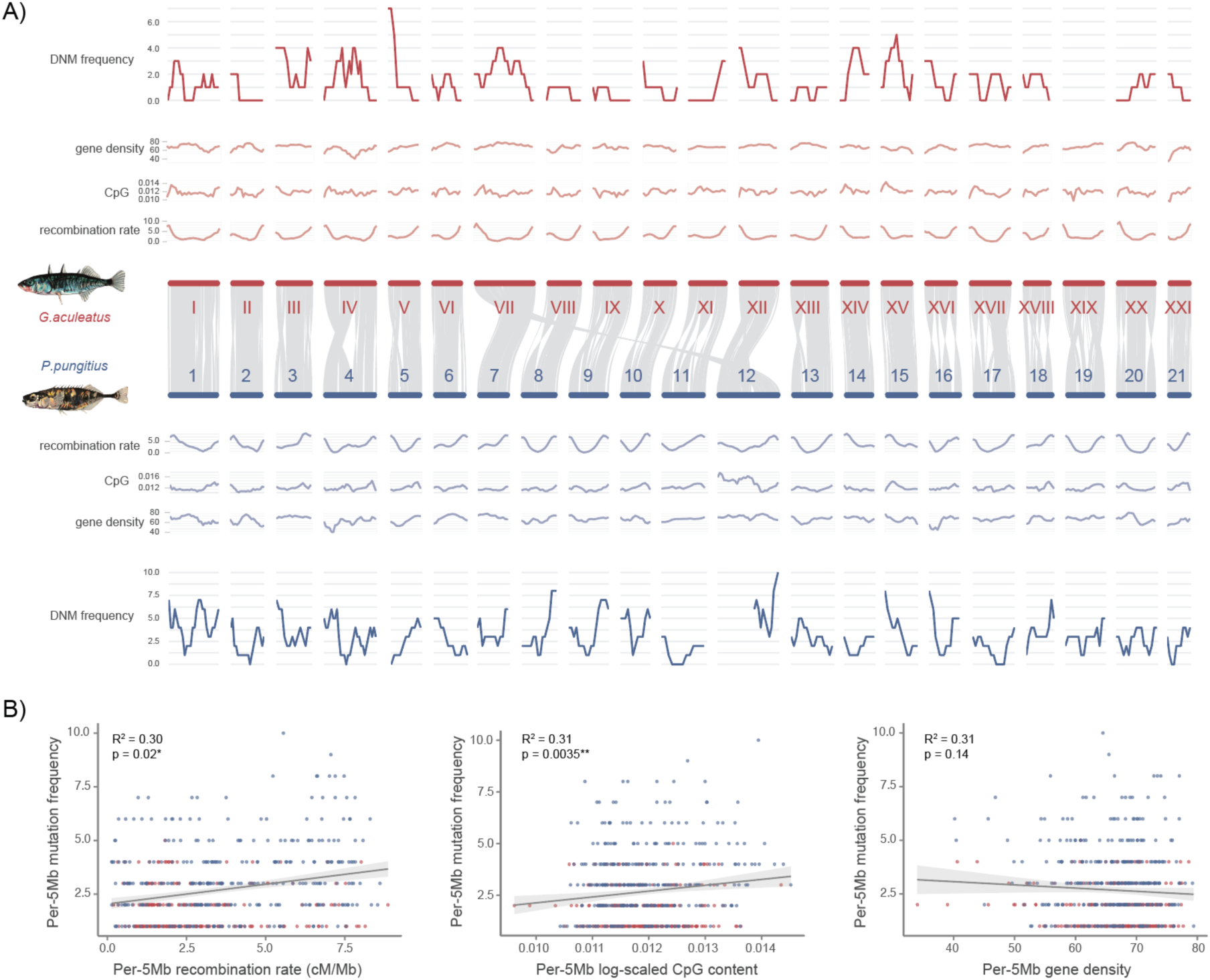
Genome-wide mutation rate and its correlation with the other genomic features. (A) The graphs, from the inner to the outer layers, displayed in order 1) the comparative synteny plot, the per-5Mb 2) recombination rates, 3) CpG content, 3) gene density and 4) DNM frequencies. (B) Correlations between the genome-wide DNM frequencies within 5Mb sliding windows and, from the left to right, 1) the recombination rates, 2) the CpG contents, and 3) the gene densities.

Furthermore, these genomic features exhibited clear correlations with each other: DNM frequencies showed a significant positive correlation with CpG contents across 5Mb sliding windows (p = 0.0035, Figure 3B). Likewise, DNM frequencies were also a positive function of recombination rate (p = 0.020, Figure 3A), but not that of gene density (p = 0.14, Figure 3B). This was also evident in the inflated *µ* in CpG sites compared to the non-CpGs (Supplementary Figure 5B), although we did not observe an obvious increase in *µ* on CpG islands (CGIs) compared to those outside CGIs, possibly due to the rarity of mutations in sticklebacks (t_16.62_ = −0.75, p = 0.46; Supplementary Figure 5A). Furthermore, a significant collinearity was observed between CpG contents and recombination rates (p = 5.74e-12, Supplementary Figure 3A). A path analysis was conducted to disentangle the direct and indirect impact of CpG content on DNM rates. The results indicated that CpG content indirectly impacted DNM rates by firstly affecting recombination rates (p < 2e-16) before impacting DNM rates (p = 0.001), while its direct impact on DNM rates was not significant (p = 0.13, Supplementary Figure 4).

## Discussion

In spite of decades of focus on the three-spined stickleback as a model system for evolutionary genetic research, no estimate of *de novo* mutation (DNM) rate has emerged for this species until now. The estimated mutation rate *µ* = 5.05 × 10^-9^/bp/gen is very similar to that obtained for *P. pungitius* (Zhang et al. 2023), and also falls in the middle of the estimates available from other teleost fishes. Furthermore, the features of the mutation spectrum and genomic characteristics of DNMs in the three-spined stickleback are very similar to those of the nine-spined stickleback suggesting little divergence in mutational processes in spite of more than 17 million years of independent evolution.

DNMs in three-spined sticklebacks were distributed randomly across the genomic features (*viz*. exonic, intronic, or intergenic), while nine-spined sticklebacks show a slight enrichment of DNMs in intergenic regions. The latter result differs slightly from that reported in Zhang et al. (2023) because of the differences in annotation files utilised. As seen in humans (Panchin et al. 2016; Youk et al. 2020) and in nine-spined sticklebacks (Zhang et al. 2023), CpG sites outside CpG islands (CGIs) exhibit higher *µ* than the genome-wide average in three-spined sticklebacks, while CpG sites within CGIs have mutation rates similar to the overall *µ*. This lower *µ* within CGIs likely contributes to the conservation and maintenance of CpG content across species. Additionally, CpG sites are known to be enriched near the transcription start sites (TSS) to regulate gene transcription (Jones 2012), and these CGIs are often hypermethylated (Youk et al. 2020). This could explain the inflated proportion of CpG DNMs in exons that was seen in both species in this study.

It is well established that mutation rates are influenced by both intrinsic and extrinsic factors. For instance, there is evidence to suggest that recombination is a mutagenic process leading to errors during DNA replication causing double-strand breaks (DSBs) (Resnick 1976; Lerciler and Hurst 2002; Halldorsson et al. 2019; Hinch et al. 2023). We found a positive correlation between the localised mutation rate and recombination rate providing evidence for the mutagenic nature of the recombination process. Highly recombining genomic regions are also enriched for CpG mutations, possibly because the methylated cytosines in single-stranded DNA are more prone to deamination (Lindahl 1993; Zhou et al. 2020).

Sticklebacks lack the functional domain of *prdm9* (Shanfelter et al. 2019; Cavassim et al. 2022), which is known to modulate the occurrence of genome-wide recombination hotspots (Baudat et al. 2010, Myers et al. 2010, Parvanov et al. 2010). As a result, recombination is more likely to occur at the promoter-like regions, including CGIs, causing the DNM rates to vary with the CpG content. Due to the stability in the distribution of GC content, the observed recombination and mutation hotspots are likely conserved for species lacking functional *prdm9* (Singhal et al. 2015). This could possibly explain why the three- and nine-spined sticklebacks exhibit similar trends in recombination rate and *µ* despite their long independent evolution. A recent study also demonstrated a similar correlation between CpG content and *µ* in dogs that have lost the entire *prdm9* (Zhang et al. 2024). The path analysis indicated that the estimated recombination rate impacted mutation rates, while being moderated by CpG content. However, given the low mutation rates in fish and the limited occurrence of DNMs within CGIs in our dataset, a more comprehensive analysis with larger sample size is needed to better understand the causality of these relationships.

The mutation spectrum observed in sticklebacks is generally similar to that of other vertebrates, with a higher prevalence of C>T and A>G substitutions compared to the other substitution types (Bergeon et al. 2023). Regardless of the presence of *prdm9*, GC-biased gene conversion (gBGC) has been found in various organisms originating from a repair bias of crossovers favouring G:C (S) over A:T (W) alleles (Birdsell 2002; Duret and Galtier 2009; Rousselle et al. 2019). This process, irrespective of its fitness effect, increases the mutation load. However, as seen in other organisms, W>S DNMs in sticklebacks are also overrepresented compared to S>W ones (Kong et al. 2012; Wong et al. 2015; Wu et al. 2020; Campbell et al. 2021; Bergeon et al. 2023). DNMs opposing gBGC lead to a decrease in the overall genomic GC content, potentially balancing the mutation load caused by gBGC. Despite these similarities, we observed a higher frequency of C>A mutations in sticklebacks compared to mammals, a trend also seen in other fish species (Bergeron et al. 2023). Given that mutation spectra are generally more similar among closely related species (Beichman et al. 2023), this phenomenon is not surprising. The elevated frequency of C>A mutations could also explain why almost all fish species exhibit transition-to-transversion (Ts:Tv) ratios at the lower end of the vertebrate spectrum (Zhang et al. 2023).

The genome-wide average DNM rates are known to be influenced both by genetic and environmental factors (Houle 1992; Drake et al. 1998). Our estimates were derived from wild-caught parents and their captive born offspring. Given that the meioses during which the detected DNMs occurred likely happened before the parents were brought to captivity, the estimated *µ* should be reflective of the natural DNM rate. To what degree the estimated rate is influenced by environmental *vs* genetic factors is not possible to deduce from our data, but whatever the relative importance of these proximate determinants is, the observed rate should be representative of the natural rate in the wild. However, whether the *µ* from this particular Baltic Sea population is representative of that from other three-spined stickleback populations from their Holarctic range remains to be verified. However, given the fairly narrow range of DNM rates in fishes (see Figure 1 in Zhang et al. 2023), it seems more likely than not that the obtained *µ* estimate is indeed representative for the species. This conjecture is also supported by the fact that the three- and nine-spined stickleback mutation rates were very similar.

Shared DNMs among full-sibs are known to occur before primordial germ cell (PGC) specification or at the early stage of post-PGC (Rahbari et al. 2016; Tang et al. 2016; Jonsson et al. 2021). The proportion of shared mutations among the three-spined stickleback full sibs was higher than that observed in mice and other fish species (Lindsay et al. 2019; Bergeron et al. 2023), but still lower than that in nine-spined sticklebacks. This could be explained by a higher number of PGCs generated in fish compared to most mammals (Lubzens et al. 2010) increasing the likelihood of early-stage mutations carried by multiple offspring (as illustrated in Supplementary Figure 2).

Species with higher percentages of shared mutations tend to have lower alpha values, indicating reduced paternal bias in DNM inheritance. Indeed, our results from three-spined sticklebacks provide further evidence for low male bias in fish, similar to findings from the nine-spined stickleback (Zhang et al. 2023) and other fish species (Bergeron et al. 2023). Additionally, we did not observe any sex bias in either the shared mutations occurring post-zygotically or the DNMs arising in the later stages. In vertebrates, no sex bias is expected during the postzygotic period before sexual differentiation of the embryos because germ cells undergo a similar number of replication events to generate spermatogonia and oogonia (Rahbari et al. 2016; Bergeron et al. 2021; 2023). However, a sex difference is expected in mammals as males typically experience more cell replication events in meiosis than females (Rahbari et al. 2016). Unlike most mammals, in which females have a limited number of genome replications of oocytes and males generate much more DNMs after puberty (Rahbari et al. 2016), female fish can produce hundreds to millions of eggs simultaneously from multiple PGCs (as detailed in Supplementary Figure 2). In addition, fish are seasonal breeders, with males producing sperm only during specific periods of the year (Bergeron et al. 2023), which further reduces the disparity in the number of reproductive cells generated by different sexes. However, lack of sex bias in *µ* could also be due to the narrow range of parental ages in our dataset, as the parents we included had just reached sexual maturity. Further tests involving a wider range of parental ages are needed to draw more definitive conclusions.

In conclusion, we have provided the first mutation rate estimate for the three-spined stickleback and found that it is very similar to that of its close relative, the nine-spined stickleback. This is somewhat surprising given that the effective population size of the three-spined stickleback is likely to be higher than that of the nine-spined stickleback (e.g. Merilä 2013; Fang et al. 2020a; Fang et al. 2021; Kemppainen et al. 2021), and because the drift-barrier hypothesis posits that species with smaller effective population sizes are expected to evolve higher mutation rates than those with larger effective sizes (Lynch 2010). Hence, one might have expected to see a higher mutation rate estimate in the three-than in the nine-spined stickleback, but this was clearly not the case. However, given the well documented differences in genomic constitutions and demographic histories across various three-spined stickleback populations (e.g. Fang et al. 2020a), future studies comparing mutation rates of freshwater and marine populations from both Pacific and Atlantic Ocean sites might be of interest.

## Acknowledgements

We thank Antoine Fraimout and Kirsi Kähkönen for access and help in obtaining the samples used in this study. Our research was supported by Seed Fund for Basic Research from University Grants Council (UGC; HKU; # 2309100132) and General Research Fund from the Research Grants Council, Hong Kong (# 17104824 to JM and KR).

## Data availability

The linkage maps applied to calculate the localised recombination rates were from Kivikoski et al. (2022, https://github.com/mikkokivikoski/InverseMappingFunctions). The raw reads of the four three-spined stickleback pedigrees used in this project will be available on ENA (accession ID: PRJEB82216), once the paper is accepted. All the lifted-over annotation files and the programming codes will be accessible on figshare at 10.6084/m9.figshare.27645810 and GitHub at https://github.com/zcharlene/3sticklebackDNMrate.

## References

Barber I. 2010. From ‘trash fish’ to supermodel: the rise and rise of the three-spined stickleback in evolution and ecology. Biologist. 7:15–21.

Bates D, Mächler M, Bolker B, Walker S. 2015. Fitting Linear Mixed-Effects Models Using lme4. J. Stat. Softw. 67(1), 1–48.

Baudat F, Buard J, Grey C, Fledel-Alon A, Ober C, Przeworski M, Coop G, de Massy B. 2010. PRDM9 is a major determinant of meiotic recombination hotspots in humans and mice. Science. 327(5967):836–40.

Beichman AC, Robinson J, Lin M, Moreno-Estrada A, Nigenda-Morales S, Harris K. 2023. Evolution of the Mutation Spectrum Across a Mammalian Phylogeny. Mol Biol Evol. 40(10):msad213.

Bell MA and Foster SA. 1994. The Evolutionary Biology of Threespine Stickleback. Oxford: Oxford University Press.

Bell AM, Bukhari SA, Sanogo YO. 2016. Natural variation in brain gene expression profiles of aggressive and nonaggressive individual sticklebacks. Behaviour. 153:1723–43

Bell AM, Trapp R, Keagy J. 2018. Parenting behaviour is highly heritable in male stickleback. R. Soc. Open Sci. 5:171029

Bergeron LA, Besenbacher S, Bakker J, Zheng J, Li P, Pacheco G, Sinding MS, Kamilari M, Gilbert MTP, Schierup MH, Zhang G. 2021. The germline mutational process in rhesus macaque and its implications for phylogenetic dating. Gigascience. 10(5):giab029.

Bergeron LA, Besenbacher S, Turner T, Versoza CJ, Wang RJ, Price AL, Armstrong E, Riera M, Carlson J, Chen HY, Hahn MW, Harris K, Kleppe AS, López-Nandam EH, Moorjani P, Pfeifer SP, Tiley GP, Yoder AD, Zhang G, Schierup MH. 2022. The mutationathon highlights the importance of reaching standardization in estimates of pedigree-based germline mutation rates. eLife. 11:e73577.

Bergeron LA, Besenbacher S, Zheng J, Li P, Bertelsen MF, Quintard B, Hoffman JI, Li Z, St Leger J, Shao C, Stiller J, Gilbert MTP, Schierup MH, Zhang G. 2023. Evolution of the germline mutation rate across vertebrates. Nature. 615(7951):285–291.

Birdsell JA. 2002. Integrating genomics, bioinformatics, and classical genetics to study the effects of recombination on genome evolution. Mol Biol Evol. 19(7):1181–97.

Campbell CR, Tiley GP, Poelstra JW, Hunnicutt KE, Larsen PA, Lee HJ, Thorne JL, Dos Reis M, Yoder AD. 2021. Pedigree-based and phylogenetic methods support surprising patterns of mutation rate and spectrum in the gray mouse lemur. Heredity (Edinb). 127(2):233–244.

Candolin U. 1997. Predation risk affects courtship and attractiveness of competing threespine stickleback males. Behav Ecol Sociobiol. 41: 81–87.

Cavassim MIA, Baker Z, Hoge C, Schierup MH, Schumer M, Przeworski M. 2022. PRDM9 losses in vertebrates are coupled to those of paralogs ZCWPW1 and ZCWPW2. Proc Natl Acad Sci U S A. 119(9):e2114401119.

Chang CC, Chow CC, Tellier LCAM, Vattikuti S, Purcell SM, Lee JJ. 2015. Second-generation PLINK: rising to the challenge of larger and richer datasets. GigaScience. 4:7.

Colosimo PF, Hosemann KE, Balabhadra S, Villarreal G Jr, Dickson M, Grimwood J, Schmutz J, Myers RM, Schluter D, Kingsley DM. 2005. Widespread parallel evolution in sticklebacks by repeated fixation of Ectodysplasin alleles. Science. 307(5717):1928–33.

Dahms C, Kemppainen P, Zanella LN, Zanella D, Carosi A, Merilä J, Momigliano P. 2022. Cast away in the Adriatic: low degree of parallel genetic differentiation in three-spined sticklebacks. Mol Ecol. 31:1234–1253.

Drake JW, Charlesworth B, Charlesworth D, Crow JF. 1998. Rates of spontaneous mutation. Genetics. 148(4):1667–86.

Duret L and Galtier N. 2009. Biased gene conversion and the evolution of mammalian genomic landscapes. Annu Rev Genomics Hum Genet. 10:285–311.

Fang B, Kemppainen P, Momigliano P, Feng X, Merilä J. 2020a. On the causes of geographically heterogeneous parallel evolution in sticklebacks. *Nat*. Ecol. Evol. 4:1105–15.

Fang B, Merilä J, Matschiner M, Momigliano P. 2020b. Estimating uncertainty in divergence times among three-spined stickleback clades using the multispecies coalescent. Mol. Phylogenet. Evol. 142:106646

Fang B, Kemppainen P, Momigliano P, Merilä J. 2021. Population structure limits parallel evolution. Mol. Biol. Evol. 38:4205–4221.

Feng X, Merilä J, Löytynoja A. 2022. Complex population history affects admixture analyses in nine-spined sticklebacks. Mol Ecol. 31:5386–5401.

Gibson G. 2005. The synthesis and evolution of a supermodel. Science. 307:1890–1891

Halldorsson BV, Palsson G, Stefansson OA, Jonsson H, Hardarson MT, Eggertsson HP, Gunnarsson B, Oddsson A, Halldorsson GH, Zink F, Gudjonsson SA, Frigge ML, Thorleifsson G, Sigurdsson A, Stacey SN, Sulem P, Masson G, Helgason A, Gudbjartsson DF, Thorsteinsdottir U, Stefansson K. 2019. Characterizing mutagenic effects of recombination through a sequence-level genetic map. Science. 363(6425):eaau1043.

Hinch R, Donnelly P, Hinch AG. 2023. Meiotic DNA breaks drive multifaceted mutagenesis in the human germ line. Science. 382(6674):eadh2531.

Houle D. 1992. Comparing evolvability and variability of quantitative traits. Genetics. 130(1):195–204.

Jones PA. 2012. Functions of DNA methylation: islands, start sites, gene bodies and beyond. Nat. Rev. Genet. 13:484–492.

Jones FC, Grabherr MG, Chan YF, Russell P, Mauceli E, Johnson J, Swofford R, Pirun M, Zody MC, White S, Birney E, Searle S, Schmutz J, Grimwood J, Dickson MC, Myers RM, Miller CT, Summers BR, Knecht AK, Brady SD, Zhang H, Pollen AA, Howes T, Amemiya C; Broad Institute Genome Sequencing Platform & Whole Genome Assembly Team; Baldwin J, Bloom T, Jaffe DB, Nicol R, Wilkinson J, Lander ES, Di Palma F, Lindblad-Toh K, Kingsley DM. 2012. The genomic basis of adaptive evolution in threespine sticklebacks. Nature. 484(7392):55–61.

Jonsson H, Magnusdottir E, Eggertsson HP, Stefansson OA, Arnadottir GA, Eiriksson O, Zink F, Helgason EA, Jonsdottir I, Gylfason A, Jonasdottir A, Jonasdottir A, Beyter D, Steingrimsdottir T, Norddahl GL, Magnusson OT, Masson G, Halldorsson BV, Thorsteinsdottir U, Helgason A, Sulem P, Gudbjartsson DF, Stefansson K. 2021. Differences between germline genomes of monozygotic twins. Nat Genet. 53(1):27–34.

Kemppainen P, Li Z, Rastas P, Löytynoja A, Fang B, Yang J, Guo B, Shikano T, Merilä J. 2021. Genetic population structure constrains local adaptation in sticklebacks. Mol Ecol. 30(9):1946–1961.

Khanna A, Larson DE, Srivatsan SN, Mosior M, Abbott TE, Kiwala S, Ley TJ, Duncavage EJ, Walter MJ, Walker JR, Griffith OL, Griffith M, Miller CA. 2022. Bam-readcount - rapid generation of basepair-resolution sequence metrics. J. Open Source Softw. 7(69), 3722

Kivikoski M, Rastas P, Löytynoja A, Merilä J. 2021. Automated improvement of stickleback reference genome assemblies with Lep-Anchor software. Mol Ecol Res. 21:2166–2176.

Kivikoski M, Rastas P, Löytynoja A, Merilä J. 2022. Predicting recombination frequency from map distance. Heredity. 130:114–121.

Koch E, Schweizer RM, Schweizer TM, Stahler DR, Smith DW, Wayne RK, Novembre J. 2019. *De novo* mutation rate estimation in wolves of known pedigree. Mol Biol Evol. 36:2536–2547.

Kong A, Frigge ML, Masson G, Besenbacher S, Sulem P, Magnusson G, Gudjonsson SA, Sigurdsson A, Jonasdottir A, Jonasdottir A, Wong WS, Sigurdsson G, Walters GB, Steinberg S, Helgason H, Thorleifsson G, Gudbjartsson DF, Helgason A, Magnusson OT, Thorsteinsdottir U, Stefansson K. 2012. Rate of *de novo* mutations and the importance of father’s age to disease risk. Nature. 488(7412):471–5.

Lescak EA, Marcotte RW, Kenney LA, von Hippel FA, Cresko WA, Sherbick ML, Colgren JJ and Andrés López J. 2015. Admixture of ancient mitochondrial lineages in three-spined stickleback populations from the North Pacific. J. Biogeogr. 42(3), pp.532–539.

Lercher MJ and Hurst LD. 2002. Human SNP variability and mutation rate are higher in regions of high recombination. Trends Genet. 18(7), 337–340.

Li H and Durbin R. 2009. Fast and accurate short read alignment with Burrows Wheeler transform. Bioinformatics. 25:1754–1760.

Li H, Handsaker B, Wysoker A, Fennell T, Ruan J, Homer N, Marth G, Abecasis G, Durbin R, 1000 Genome Project Data Processing Subgroup. 2009. The sequence alignment/map (SAM) format and SAMtools. Bioinformatics. 25:2078–2079

Lindahl T. 1993. Instability and decay of the primary structure of DNA. Nature. 362(6422):709–15.

Lindsay SJ, Rahbari R, Kaplanis J, Keane T, Hurles ME. 2019. Similarities and differences in patterns of germline mutation between mice and humans. Nat Commun. 10:4053.

Liu S, Ferchaud AL, Grønkjaer P, Nygaard R, Hansen MM. 2018. Genomic parallelism and lack thereof in contrasting systems of three-spined sticklebacks. Mol Ecol. 27:4725–4743.

Lopez F, Charbonnier G, Kermezli Y, Belhocine M, Ferré Q, Zweig N, Aribi M, Gonzalez A, Spicuglia S, Puthier D. 2019. Explore, edit and leverage genomic annotations using Python GTF toolkit. Bioinformatics. 35(18):3487–3488.

Lubzens E, Young G, Bobe J, Cerdà J. 2010. Oogenesis in teleosts: how eggs are formed. Gen Comp Endocrinol. 165(3):367–89.

Lynch M. 2010. Evolution of the mutation rate. Trends Genet. 26:345–352.

Mäkinen HS, Cano JM, Merilä J. 2006. Genetic relationships among marine and freshwater populations of the European three-spined stickleback (*Gasterosteus aculeatus*) revealed by microsatellites. Mol Ecol. 15(6):1519–34.

Mäkinen HS and Merilä J. 2008. Mitochondrial DNA phylogeography of the three-spined stickleback (*Gasterosteus aculeatus*) in Europe-evidence for multiple glacial refugia. Mol Phylogenet Evol. 46(1):167–82.

Merilä J. 2013. Nine-spined stickleback (*Pungitius pungitius*): an emerging model for evolutionary biology research. Ann N Y Acad Sci. 1289:18–35.

Miklem G and Hillier L. Unpublished data:http://genome.ucsc.edu/cgi-bin/hgTrackUi?g=cpgIslandExt

Myers S, Bowden R, Tumian A, Bontrop RE, Freeman C, MacFie TS, McVean G, Donnelly P. 2010. Drive against hotspot motifs in primates implicates the PRDM9 gene in meiotic recombination. Science. 327(5967):876–9.

Norton WHJ and Gutiérrez HC. 2019. The three-spined stickleback as a model for behavioural neuroscience. PLoS One. 14(3),e0213320.

Ortí G, Bell MA, Reimchen TE, Meyer A. 1994. Global survey of mitochondrial DNA sequences in the threespine stickleback: evidence for recent migrations. Evolution. 48(3): 608–622.

Panchin AY, Makeev VJ, Medvedeva YA. 2016. Preservation of methylated CpG dinucleotides in human CpG islands. Biol Direct. 11(1):11.

Parvanov ED, Petkov PM, Paigen K. 2010. Prdm9 controls activation of mammalian recombination hotspots. Science. 327(5967):835.

Peichel CL and Marques DA, 2017. The genetic and molecular architecture of phenotypic diversity in sticklebacks. Philos. Trans. R. Soc. Lond. B Biol. Sci. 372(1713), p20150486

Peichel CL, McCann SR, Ross JA, Naftaly AFS, Urton JR, Cech JN, Grimwood J, Schmutz J, Myers RM, Kingsley DM, White MA. 2020. Assembly of the threespine stickleback Y chromosome reveals convergent signatures of sex chromosome evolution. Genome Biol. 21(1):177.

Poplin R, Ruano-Rubio V, DePristo MA, Fennell TJ, Carneiro MO, Van der Auwera GA, Kling DE, Gauthier LD, Levy-Moonshine A, Roazen D, Shakir K, Thibault J, Chandran S, Whelan C, Lek M, Gabriel S, Daly MJ, Neale B, MacArthur DG, Banks E. 2017. Unpublished data: https://www.biorxiv.org/content/10.1101/201178v3.

Rahbari R, Wuster A, Lindsay SJ, Hardwick RJ, Alexandrov LB, Turki SA, Dominiczak A, Morris A, Porteous D, Smith B, Stratton MR; UK10K Consortium; Hurles ME. 2016. Timing, rates and spectra of human germline mutation. Nat Genet. 48(2):126–133.

Ravinet M, Yoshida K, Shigenobu S, Toyoda A, Fujiyama A, Kitano J. 2018. The genomic landscape at a late stage of stickleback speciation: high genomic divergence interspersed by small localized regions of introgression. PLoS Genet. 14:e1007358.

Reid K, Bell MA, Veeramah KR. 2021. Threespine stickleback: A model system for evolutionary genomics. Annu. Rev. Genomics Hum. Genet. 22:357–383.

Resnick MA. 1976. The repair of double-strand breaks in DNA; a model involving recombination. J Theor Biol.; 59(1):97–106.

Roberts Kingman GA, Vyas DN, Jones FC, Brady SD, Chen HI, Reid K, Milhaven M, Bertino TS, Aguirre WE, Heins DC, von Hippel FA, Park PJ, Kirch M, Absher DM, Myers RM, Di Palma F, Bell MA, Kingsley DM, Veeramah KR. 2021. Predicting future from past: The genomic basis of recurrent and rapid stickleback evolution. Sci Adv.7(25):eabg5285.

Rousselle M, Laverré A, Figuet E, Nabholz B, Galtier N. 2019. Influence of Recombination and GC-biased Gene Conversion on the Adaptive and Nonadaptive Substitution Rate in Mammals versus Birds. Mol Biol Evol. 36(3):458–471.

Sendell-Price AT, Tulenko FJ, Pettersson M, Kang D, Montandon M, Winkler S, Kulb K, Naylor GP, Phillippy A, Fedrigo O, Mountcastle J. 2023. Low mutation rate in epaulette sharks is consistent with a slow rate of evolution in sharks. Nat. Commun., 14(1), 6628

Shanfelter AF, Archambeault SL, White MA. 2019. Divergent fine-scale recombination landscapes between a freshwater and marine population of threespine stickleback fish. Genome biol. evol. 11(6), 1552–1572.

Shikano T, Shimada Y, Herczeg G, Merilä J. 2010. History vs. habitat type: explaining the genetic structure of European nine-spined stickleback (*Pungitius pungitius*) populations. Mol Ecol. 19(6):1147–61.

Singhal S, Leffler EM, Sannareddy K, Turner I, Venn O, Hooper DM, Strand AI, Li Q, Raney B, Balakrishnan CN, Griffith SC, McVean G, Przeworski M. 2015. Stable recombination hotspots in birds. Science. 350(6263):928–32.

Smeds L, Qvarnström A, Ellegren H. 2016. Direct estimate of the rate of germline mutation in a bird. Genome Res. 26:1211–1218.

Tang H, Krishnakumar V, Zeng X, Xu Z, Taranto A, Lomas JS, Zhang Y, Huang Y, Wang Y, Yim WC, Zhang J, Zhang X. 2024. JCVI: A versatile toolkit for comparative genomics analysis. Imeta. 3(4):e211.

Tang WW, Kobayashi T, Irie N, Dietmann S, Surani MA. 2016. Specification and epigenetic programming of the human germ line. Nat Rev Genet. 17(10):585–600.

Thorvaldsdóttir H, Robinson JT, Mesirov JP. 2013. Integrative genomics viewer (IGV): high-performance genomics data visualization and exploration. Brief Bioinform. 14:178–192.

Van der Auwera GA and O’Connor BD. 2020. Genomics in the cloud: using docker, GATK, and WDL in Terra (1st edition). Sebastopol: O’Reilly Media.

Varadharajan S, Rastas P, Löytynoja A, Matschiner M, Calboli FCF, Guo B, Nederbragt AJ, Jakobsen KS, Merilä J. 2019. A high-quality assembly of the nine-spined stickleback (*Pungitius pungitius*) genome. Geno Biol Evol. 11:3291–3308.

Von Hippel F. 2010. Tinbergen’s legacy in behaviour: sixty years of landmark stickleback papers. Leiden: Brill.

Von Hippel FA, Trammell EJ, Merilä J, Sanders MB, Schwarz T, Postlethwait JH, Titus TA, Buck CL, Katsiadaki I. 2016. The ninespine stickleback as a model organism in arctic ecotoxicology. Evol. Ecol. Res. 17: 487–504

Wang RJ, Raveendran M, Harris RA, Murphy WJ, Lyons LA, Rogers J, Hahn MW. 2022. *De novo* Mutations in Domestic Cat are Consistent with an Effect of Reproductive Longevity on Both the Rate and Spectrum of Mutations. Mol Biol Evol. 39(7):msac147.

Wang Y and Obbard DJ, 2023. Experimental estimates of germline mutation rate in eukaryotes: a phylogenetic meta-analysis. Evol. Lett. 7(4),216–226.

Wootton RJ. 2009. The Darwinian stickleback *Gasterosteus aculeatus*: a history of evolutionary studies. J. Fish Biol. 75(8), 1919–1942.

Wong WS, Solomon BD, Bodian DL, Kothiyal P, Eley G, Huddleston KC, Baker R, Thach DC, Iyer RK, Vockley JG, Niederhuber JE. 2016. New observations on maternal age effect on germline de novo mutations. Nat Commun. 7:10486.

Wu FL, Strand AI, Cox LA, Ober C, Wall JD, Moorjani P, Przeworski M. 2020. A comparison of humans and baboons suggests germline mutation rates do not track cell divisions. PLoS Biol. 18(8):e3000838.

Yamasaki YY, Kakioka R, Takahashi H, Toyoda A, Nagano AJ, Machida Y, Møller PR, Kitano J. 2020. Genome-wide patterns of divergence and introgression after secondary contact between *Pungitius* sticklebacks. Philos T Roy Soc B. 375:20190548.

Yoder AD and Tiley GP. 2021. The challenge and promise of estimating the de novo mutation rate from whole-genome comparisons among closely related individuals. Mol Ecol. 30(23):6087–6100.

Youk J, An Y, Park S, Lee JK, Ju YS. 2020. The genome-wide landscape of C:G > T:a polymorphism at the CpG contexts in the human population. BMC Genomics. 21:270.

Zhang C, Reid K, Sands AF, Fraimout A, Schierup MH, Merilä J. 2023. *De Novo* Mutation Rates in Sticklebacks. Mol Biol Evol. 40(9):msad192.

Zhang SJ, Ma J, Riera M, Besenbacher S, Salokorpi N, Hundi S, Hytönen MK, Zhou T, Ostrander EA, Schierup MH, Lohi H, Wang GD. 2024. Unpublished data: 10.1101/2024.06.04.596747.

Zhou Y, He F, Pu W, Gu X, Wang J, Su Z. 2020. The Impact of DNA Methylation Dynamics on the Mutation Rate During Human Germline Development. G3 (Bethesda). 10(9):3337–3346.

