## Supplementary for "Rate of *de novo* mutations in the three-spined stickleback"

*Supplementary Figure 1.* Comparisons of standardised number of DNMs inherited from the two parents on CpG sites (with light grey border) and non-CpG sites (with dark grey border).

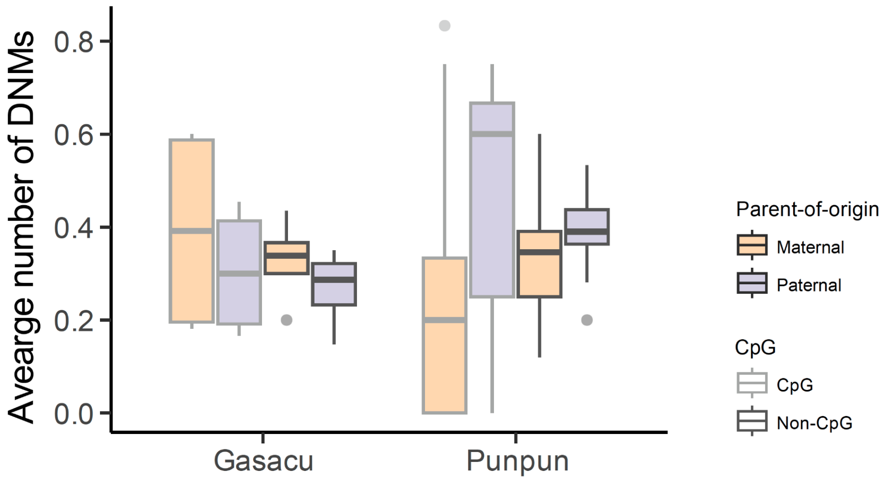

*Supplementary Figure 2.* A detailed illustration of embryogenesis and gametogenesis in sticklebacks. Spermatogenesis on the left (purple), oogenesis on the right (orange), and the corresponding life stages in the middle (blue). Maternal inheritance of 'nuage' determines the functions of cells after the 1000-cell stage in fish, including germ cells that migrate to the gonads before maturation (represented as yellow cells in the middle life-stage diagram). This process is known as primordial germ cell specification (PGCS). If mutations arise before PGCS or early in the post-PGCS stage (as depicted on the right side of the maternal germline), these mutations will be present in following-stage germ cells and can be shared among siblings. Conversely, if the mutation occurs later during meiosis (as shown on the left side of the paternal germline), the likelihood of this DNM being shared among siblings is little.

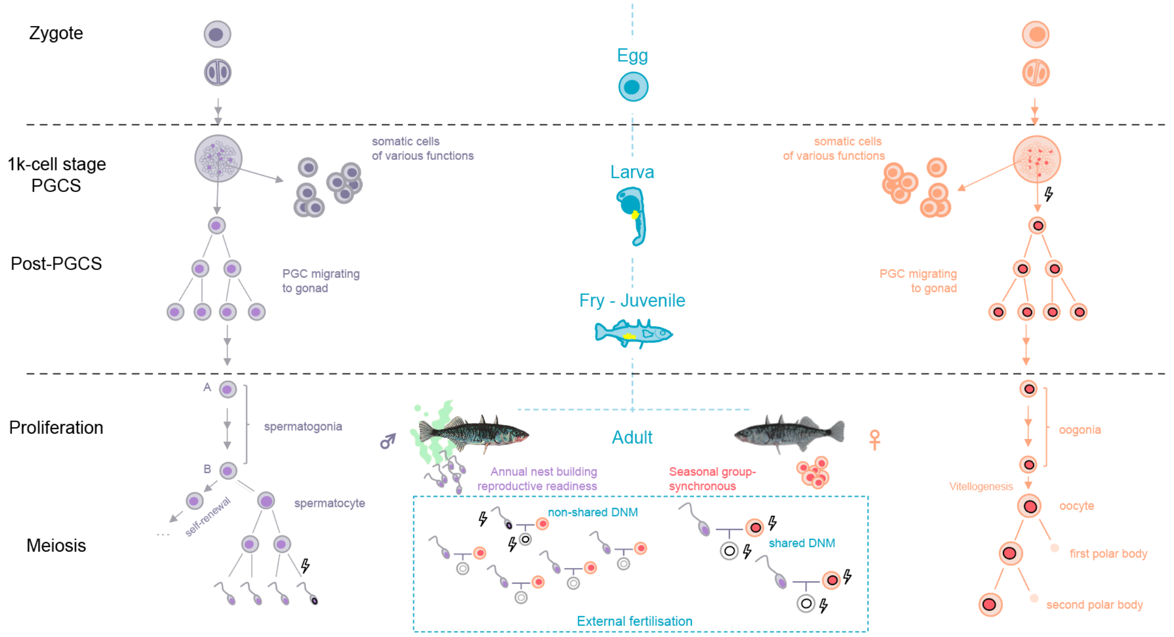

*Supplementary Figure 3.* (A) A correlation between the per-5Mb CpG contents and the recombination rates. (B) Correlation of the per-chromosomal mutation rates between the two species, accounting for their synteny.

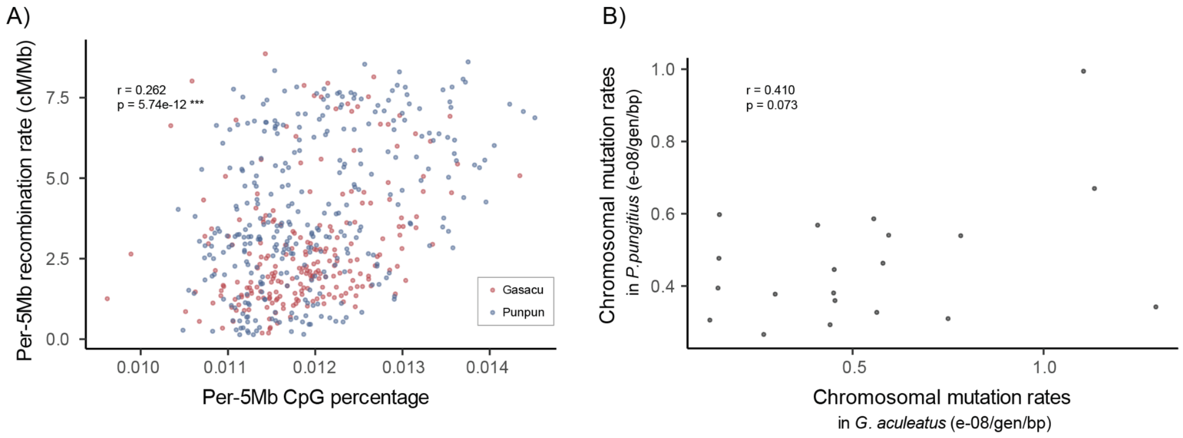

*Supplementary Figure 4.* A path analysis of the direct and indirect impacts of CpG content and recombination rate on DNM rates.

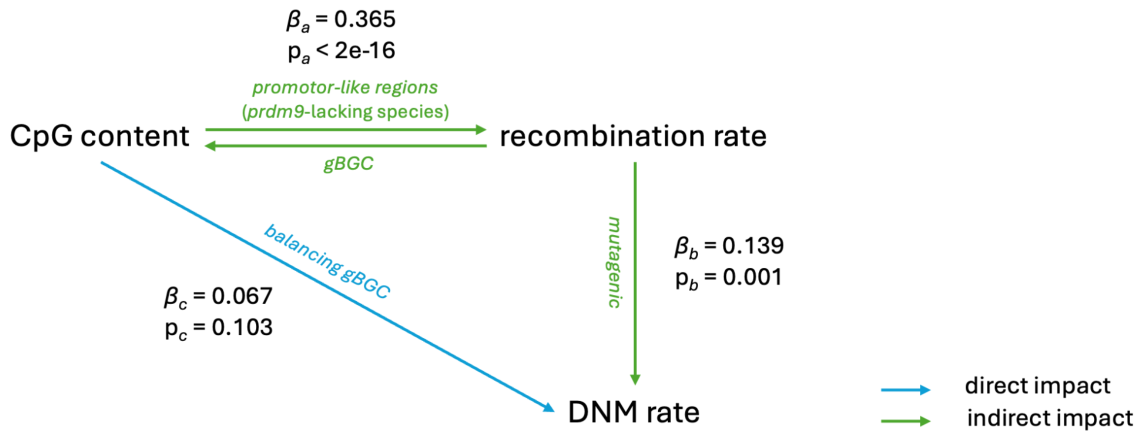


*Supplementary Figure 5.* Rates of de novo mutations (/bp/generation) compared between A) CG sites located within and outside CpG island, and B) CpG sites and non-CpG sites.

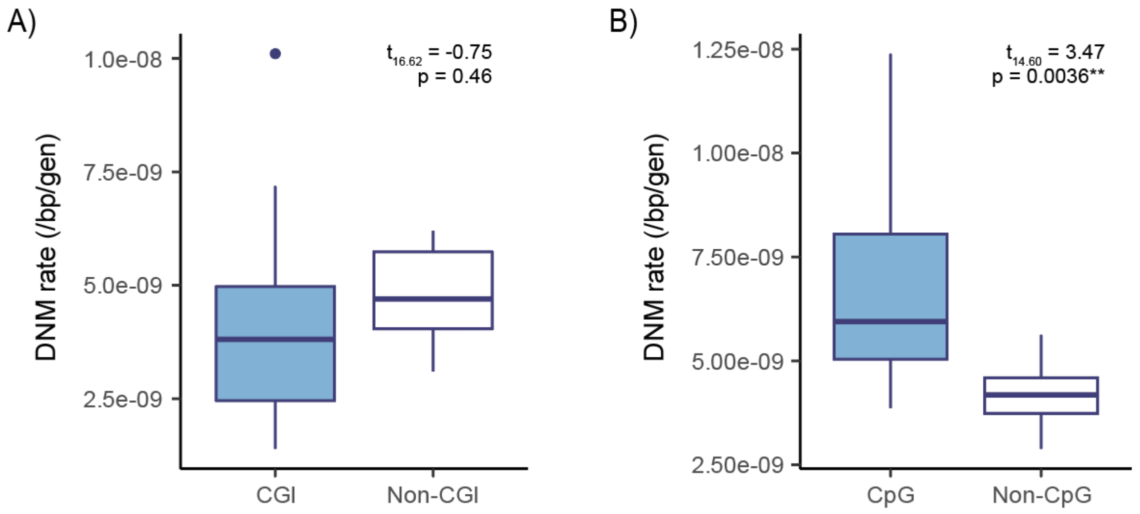

*Supplementary Table 1.* Filters applied identifying de novo mutations (DNM) and estimating the DNM rates.

| **Filter** | **Detail** |
|---|---|
| Site filters | Quality by depth (QD) < 2.0<br>Mapping quality (MQ) < 40.0<br>Fisher's exact test on strand bias (FS) > 60.0<br>Strand odds ratio (SOR) > 3.0<br>Mapping quality rank sum test (MQRankSum) < −12.5<br>Read position rank sum test (ReadPosRankSum) < −8.0 |
| Individual filters | sequencing depth (DP ≤ 20 and DP ≥ 100)<br>genotyping quality (GQ ≤ 80)<br>allelic depth filter (AD1 > 0 for parents)<br>allelic balance (AB < 0.3 and AB > 0.7 for offspring)<br>sequencing depth (DP < $0.5DP_{trio}$ and DP > $2DP_{trio}$ for offspring)<br>DNM candidates within 5 bp away from indels<br>DNM candidates that occurred in unrelated samples |
| Post-filtering check (False positives) | IGV manual curation and bam-readcount (excluding sites where the parents carry alternative alleles or the offspring do not have a sufficient number of alternative alleles) |
| Callable genome size | Number of sites passing the trio depth filtering ($0.5DP_{trio} < DP_{child} < 2DP_{trio}$) |
| False negatives | Number of true heterozygotes deleted by the allelic balance filtering (AB < 0.3 and AB > 0.7) |
